# The *Lotus japonicus* alpha Expansin *EXPA1* is recruited during intracellular and intercellular rhizobial colonization

**DOI:** 10.1101/2025.09.08.674960

**Authors:** Jesús Montiel, Ivette García Soto, Elizabeth Monroy Morales, Beatrice Lace, Mads Vestergaard, Niels Sandal, Thomas Ott, Jens Stougaard

**Author notes:** Correspondence: Jesús Montiel. Correspondence: Jens Stougaard. **Significance statement:** The colonization of legume roots by rhizobia represents a critical phase in the establishment of root nodule symbiosis. In this study, we demonstrated that LjEXPA1, a plant protein involved in cell wall loosening, is indispensable in the two modalities of rhizobial symbiotic infection in *Lotus japonicus* roots.

## Abstract

Most legumes establish a mutualistic association with rhizobia, a group of nitrogen-fixing bacteria. In *Lotus japonicus*, the symbiotic colonization occurs intracellularly, via root-hair infection threads by *Mesorhizobium loti*, or intercellularly, with *Agrobacterium pusense* IRBG74. In both mechanisms, cell wall remodeling is presumably an essential process. In plants, alpha-Expansins (EXPA) promote cell wall loosening by non-enzymatically triggering a pH-dependent relaxation. In this study, we show that LjEXPA1, is critical for the intracellular and intercellular symbiotic program in *L. japonicus*. Promoter activity and subcellular localization analyses revealed that EXPA1 is recruited at essential compartments and structures of epidermal and cortical cells in both mechanisms of rhizobial infection, such as the infection chambers, infection pockets and transcellular infection threads. Additionally, EXPA1-YFP abundantly accumulated in dividing cortical cells during nodule formation. The expression profile of *EXPA1* correlates with the symbiotic phenotype observed in homozygous mutants disrupted in the *EXPA*1 gene (*expA1*-1 and *expA1*-2). Infection thread formation and intercellular colonization was drastically reduced in *expA1*-1 and *expA1*-2 mutants, respect to wild type plants. Similarly, nodule formation was significantly reduced in these mutants after *M. loti* or IRBG74 inoculation. Our results indicate that non-enzymatic cell wall remodeling by the alpha Expansin EXPA1 is crucial for the successful establishment of *Lotus*-rhizobia symbiosis, regardless of the infection mechanism.

## INTRODUCTION

Nitrogen is a limiting macronutrient for plant growth; however, legumes can obtain it through an endosymbiotic relationship with nitrogen-fixing soil bacteria, known as rhizobia. This legume-rhizobial symbiosis leads to the formation of root-derived organs known as nodules (Downie, 2014). A successful symbiosis between legumes and rhizobia is initiated by a molecular dialogue, which requires two coordinated developmental processes: the bacterial infection in the epidermis, which allows bacteria to invade host cells, and the induction of cortical cell divisions, leading to the formation of nodule primordia (Oldroyd *et al*., 2011, Lace and Ott, 2018). Although infection and organogenesis are highly synchronized, these are genetically separable processes. Nodulation requires host perception of lipochitooligosaccharide signals, known as Nod factors, which are synthesized by rhizobia in response to plant-derived flavonoids. In *Lotus japonicus*, Nod factors are recognized by two LysM receptor-like kinases, Nod Factor Receptor 1 (*Lj*NFR1) and *Lj*NFR5, which trigger a downstream signaling cascade (Radutoiu *et al*., 2003, Bozsoki *et al*., 2020, Rubsam *et al*., 2023).

Rhizobial infection in legumes can occur via two primary routes: transcellular (infection thread (IT)-mediated) and intercellular (without epidermal infection threads). Intracellular entry is the best-known infection process particularly in model legumes like *Medicago truncatula* and *L. japonicus* (de Carvalho-Niebel *et al*., 2024). In this process, rhizobia attach to elongating root hairs, inducing growth redirection and entrapment within a curl, forming an infection chamber (IC). From this chamber, the cell wall and plasma membrane invaginate to form an inward, transcellular tunnel-like structure, known as IT, to mediate the entry of rhizobia and guide their proliferation across root tissues (Gao *et al*., 2024). Throughout the progression of rhizobial infection, organogenesis is executed in root cortical and pericycle cells, resulting in the development of a nodule primordium. As the infection threads transcellularly grow, they penetrate several cortical cell layers and release bacteria inside the nodule primordia cells (Monroy-Morales *et al*., 2022). These colonized nodule cells create a low-oxygen environment that supports the activity of the nitrogenase complex in bacteroids, a bacterial enzyme that reduces atmospheric nitrogen into ammonia, which plants can then assimilate through a process known as biological nitrogen fixation (Ferguson *et al*., 2010).

Although the transcellular infection pathway is well-documented, approximately 25% of legume species undergo intercellular infection (Sprent *et al*., 2017). In this infection mode, rhizobia enter the root through sites of epidermal cracking or fissures, followed by intercellular penetration. Species such as *Sesbania rostrata*, *Arachis hypogaea* (peanut), *Lupinus albus* and *Lotus uliginosus,* exhibit nodules infected without epidermal ITs (Ibañez *et al*., 2017). Furthermore, *L. japonicus* can also be infected through both transcellular and intercellular routes (Quilbe *et al*., 2022). While *L. japonicus* typically undergoes transcellular infection via ITs, some plant mutants lacking epidermal ITs can still form nodules and internalize rhizobia, albeit at a low frequency (Karas *et al*., 2005, Madsen *et al*., 2010). Additionally, certain strains of rhizobia, such as *Sinorhizobium fredii* HH103, *Rhizobium leguminosarum* Norway, and *Agrobacterium pusense* IRGB74, colonized *Lotus* roots intercellularly (Acosta-Jurado *et al*., 2016, Liang *et al*., 2019, Montiel *et al*., 2021, Zarrabian *et al*., 2022). This highlights the flexibility of *L. japonicus* in accommodating both infection routes depending on the rhizobial strain and environmental conditions.

A key aspect of rhizobial infection is cell wall remodeling, which is essential for allowing rhizobia to enter root cells and for the reactivation of meristematic activity during nodule development (Rich *et al*., 2014). The plant cell wall is a dynamic structure, mainly composed by pectin, hemicellulose and cellulose, that can be altered in structure and composition by the action of a complex and sophisticated network of numerous protein families (Broxterman and Schols, 2018). Nonetheless, the evidence collected by different research groups indicates that certain members of these cell wall protein families have a pivotal role in the legume-rhizobia symbiosis. In *L. japonicus*, nodule development is negatively affected in the *cellulose synthase-like d1* mutant (Karas *et al*., 2021), while in *M. truncatula* the *Glycoside Hydrolase 9C2 (GH9C2)* gene is required for both epidermal and nodule infection (Zhao *et al*., 2025). Similarly, a nodulation pectate lyase gene (*NPL*) is induced in the nodulation process of *L. japonicus* after *M. loti* inoculation, and the *Ljnpl* mutant exhibit empty nodules and abnormal root-hair ITs (Xie *et al*., 2012). In *M. truncatula* root hairs, the coordinated action of a symbiosis-specific pectin methyl esterase (SyPME1) and MtNPL, allows intracellular progression of *Sinorhizobium meliloti* (Su *et al*., 2023) and MtNPL was also shown to act synergistically with GH9C2 on rhizobial infection (Zhao *et al*., 2025). *In vivo* localization of cell wall proteins, such as ENOD11, NPL, SyPME1 and GH9C2, has been observed at early infection sites, supporting cell wall remodeling in radially-expanding chambers to facilitate IT growth (de Carvalho-Niebel *et al*., 2024). Remodeling may occur at the transcellular passage cleft (TPC), supporting cell-to-cell progression of the IT and resulting in the release of bacteria into dividing primordium cells (Roy *et al*., 2020, Zhang and Ott, 2024).

The mechanical properties of the cell wall can be modified by the non-enzymatic action of Expansins, a large family of extracellular proteins with a carbohydrate binding module (Cosgrove, 2015). Expansins are classified into four subfamilies based on phylogenetic analysis and sequence characteristics: α-Expansin (EXPA), β-Expansin (EXPB), Expansin-like A (EXLA), and Expansin-like B (EXLB). Both α-expansins and β-expansins are well-known for their ability to relax the cell wall, aiding in processes like cell expansion during growth and development (Cosgrove, 2015). In the context of symbiosis, the overexpression of the soybean β-expansin *GmEXPB2*, enhances root growth, increases rhizobial infection events, promotes primordia formation, and boosts the number of mature nodules (Li *et al*., 2015). However, the transcriptome analysis in different legumes revealed that during nodulation, the expression of several isoforms of the alpha Expansin subfamily are significantly induced (Gyorgyey *et al*., 2000, Giordano and Hirsch, 2004, Libourel *et al*., 2023). Herein to get further insights into the role of alpha Expansins in the Legume-rhizobia symbiosis, we analyzed *LjEXPA1* during the symbiotic program induced by *M. loti* and *A. pusense* IRBG74, that colonize *L. japonicus* roots, intra- and intercellularly, respectively.

## MATERIALS AND METHODS

### Biological Materials and Nodulation Kinetics

In this study, wild type and mutant lines belong to the genetic background of *L. japonicus* accession Gifu (Handberg and Stougaard, 1992). The mutant lines 30015381 and 30164465 were obtained from the *LORE1* collection and genotyped, as described in the database, to obtain homozygous mutants (Malolepszy *et al*., 2016). The seeds were scarified with sandpaper, sterilized with a 5% chlorine solution for 15 minutes and washed with sterile distilled water (3-5 times) to remove any residual chlorine. The disinfected seeds were germinated in 12 cm x 12 cm square Petri dishes containing 1.2% Bacto agar in a growth chamber at 21°C. Seedlings at 3-7 days post-germination (dpg) were transferred to new square Petri dishes containing ¼ B&D solid medium (Broughton and Dilworth, 1971) and filter paper, and then placed in a growth room with photoperiod (16:8 at 21°C). For nodulation kinetics, the seedlings were inoculated with 1 mL of *M. loti* R7A::DsRed, *M. loti* R7A-GFP or IRBG74::DsRed (OD 600nm= 0.05), previously grown in YEM medium. Nodule numbers were recorded at 1-6 weeks post-inoculation (wpi) using a stereomicroscope.

### Phylogenetic Analysis

To identify members of the Expansin superfamily in *Lotus*, BLASTP analyses were performed using as queries the 42 annotated Expansin protein sequences from *M. truncatula*. Homologous protein sequences were also retrieved from the Expansin Gene Family Database (http://www.expansingenefamily.com) for the following species: *Glycine soja* (*G. soja*) (Feng *et al*., 2022), *Arabidopsis thaliana* (*A. thaliana*) (https://www.arabidopsis.org), *Physcomitrium patens* (*P. patens*), *Oryza sativa* (*O. sativa*), *Sorghum bicolor* (*S. bicolor*), Solanum pennellii (S. pennellii), *Capsella rubella* (*C. rubella*), and *Phaseolus vulgaris* (*P. vulgaris*) (Kok *et al*., 2023). Protein sequences from *Lotus* and the ten additional species were aligned using ClustalX2 (Larkin *et al*., 2007), and a phylogenetic tree was inferred by the Neighbor-Joining method. Tree topology was evaluated, and the graphical representation of the tree was edited and exported using the Interactive Tree of Life (iTOL) web interface (https://itol.embl.de)

### Intercellular and intracellular rhizobial infection

Gifu and *expA* mutant lines were inoculated with the *M. loti*–LacZ or IRBG74, as described for nodulation kinetics. For intracellular infection, roots were collected at 10 dpi with *M. loti* and stained with a 2 mg/mL X-Gal solution to visualize and count the epidermal and cortical ITs under an optical microscope. The intercellular infection was analyzed as previously described by Montiel et al. (2021). Gifu and mutant roots were collected at 3 weeks post-inoculation (wpi) with IRBG74, incubated for 1 min in a solution for surface disinfection (0.3% *w*/*v* of hypochlorite and 70% *v*/*v* EtOH), and then washed 5 times with distilled water. The total DNA was extracted from individual roots, adjusted to a final concentration of 10 ng µL−1, and used as a template for qPCR to evaluate the IRBG74 *NodA* abundance with the primers; forward: GAACTGCAAGTTGACGATCACGC and reverse: AAACGTCGTAACAAGCCCATGTGG. The expression values were normalized to the abundance of the *L. japonicus* gene LotjaGi1g1v0152000.1 with the oligonucleotides; forward: GAAGGACCCAGAGGATCACA and reverse: CGGTCTTCGTACTTCTTCGC using the delta Ct method (Pfaffl, 2001).

### Constructs for promoter activity and subcellular localization

For *pLjEXPA1::tYFP-nls*, and *pLjEXPA1::LjEXPA1-YFP_p35S::DsRed-nls* constructs, the predicted promoter sequence (2 kb upstream of the start codon) and coding sequence (CDS) were obtained from the *L. japonicus* Gifu genome in the *Lotus* database. Each module was assembled using Golden Gate technology, and the resulting constructs were used to transform *Agrobacterium rhizogenes* strain AR1193. Subsequently, Gifu roots were transformed according to a standardized protocol (Hansen *et al*., 1989). Transgenic roots were inoculated with *M. loti* R7A-DsRed (Kelly *et al*., 2013) or IRBG74-DsRed (Montiel *et al*., 2021), and examined by confocal microscopy. For subcellular localization imaging, bacterial strains were inoculated using an OD600 = 0.01 (*M. loti* R7A-DsRed) or OD600 = 0.005 (IRBG74-DsRed).

### Confocal Microscopy Analysis

The first set of observations, focused on promoter activity, was made using an inverted confocal laser scanning microscope (FV1000) equipped with a 40×/NA 0.75 dry objective. YFP and DsRed signals were detected using excitation wavelengths of 488 nm and 543 nm, respectively, with emission windows set at 505-525 nm and 560-660 nm to collect their signal.

Subcellular localization imaging was performed using a Leica SP8 FALCON FLIM confocal microscope equipped with a 20 x/0.75 water immersion lens (Leica Microsystems, Mannheim, Germany). A pulsed White Light Laser (WLL) was used as an excitation source. YFP and DsRed were excited at 513 nm and 561 nm, respectively, and their emissions were collected at 520–550 nm and 580–620 nm. Before inoculating the bacterial strains, the position of the apex of transformed roots was marked on plates. At 4–5 dpi (M. loti R7A-DsRed) or 8-12 dpi (IRBG74-DsRed), root segments that had grown beyond the mark were excised, mounted in water, and imaged.

### Statistical analyses

All statistical analyses were performed using GraphPad Prism 10, and graphical representations were made using R. Individual comparisons were done by Mann–Whitney *U* test. *P*-values and the number of samples are shown in the figure legends.

## RESULTS

### Two *Lotus* alpha *Expansins* are upregulated by *M. loti* and IRBG74

To investigate the potential role of *Lotus* Expansins in the legume-rhizobia symbiosis, we first performed a search in the *Lotus* genome for homologous genes to soybean and *Arabidopsis thaliana Expansins*, detecting 37 genes (Table S1). To narrow down the set of *Lotus* Expansins that are likely recruited during the symbiotic process, the expression profile of these sequences was explored in the Lotus Expression Atlas (Mun *et al*., 2016, Kamal *et al*., 2020). This analysis revealed that a relatively low number of Expansins are induced during root nodule symbiosis (RNS), or in root hairs, where intracellular rhizobial colonization occurs (Figure S1). To identify which isoforms might participate in the intracellular and intercellular infection program, a gene expression heatmap was made with the available RNA-seq data of *Lotus* roots inoculated with *M. loti* and IRBG74 (Montiel *et al*., 2021). Only three Expansin were significantly induced by both symbionts at different timepoints: LotjaGi1g1v0089200, LotjaGi2g1v0429600 and LotjaGi1g1v0606600 (Figure 1A).

**Figure 1.**
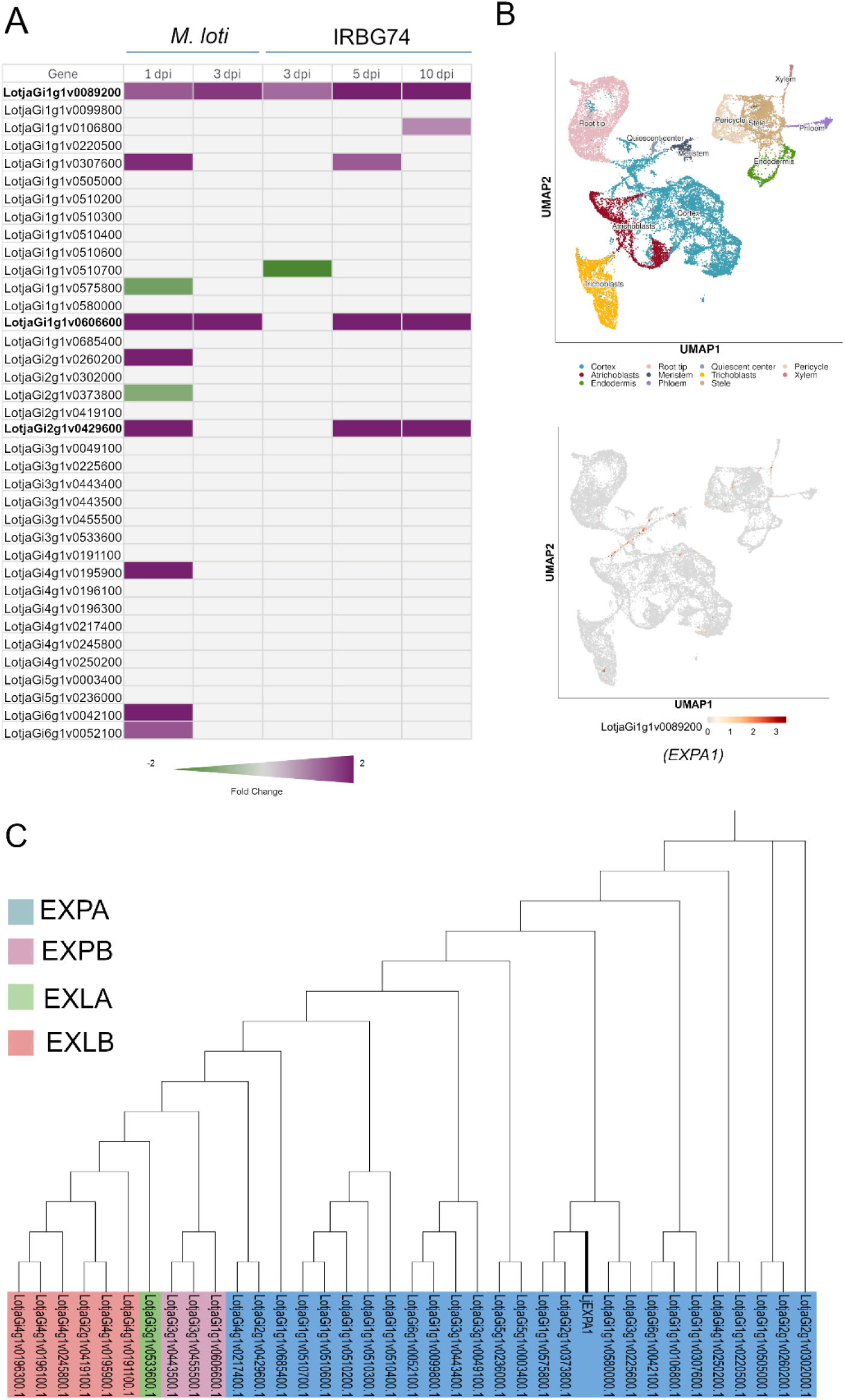
Expression profile and phylogeny of *Lotus* Expansins. **A,** Heat map expression of *Lotus* genes encoding Expansins in roots inoculated with *M. loti* and IRBG74 (extracted from RNAseq data; Montiel et al. 2021). **B,** Expression pattern of *LjExpA1* (LotjaGi1g1v0089200) extracted from single-cell RNAseq data (Frank et al. 2023). **C,** Phylogenetic distribution of *Lotus* Expansins across four subfamilies: alpha, beta, alpha-like, and beta-like.

To obtain additional information of *Lotus* Expansins, the amino acid sequences for the 37 genes were predicted, and a phylogenetic tree was constructed. This analysis included the Expansin family of representative legumes (*Glycine soja*, *Phaseolus vulgaris* and *Medicago truncatula*) and non-legumes (*Oryza sativa* and *A. thaliana*) plant species. The resulting phylogenetic tree showed that for all species tested, the EXPA subfamily is the most abundant, followed by EXLB (Figure S2A and B). The phylogenetic distribution of the four *Lotus* Expansin subfamilies was also consistent with their clustering by an identity matrix (Figure S2C). In addition, this analysis revealed that one of the *Expansin* genes induced by *M. loti* and IRBG74, encodes an Alpha subfamily isoform (Figure 1C), hereafter referred as *EXPA1* (LotjaGi1g1v0089200). Interestingly, in the single cell RNA-seq data of *L. japonicus* roots at 10 days post-inoculation (dpi) with *M. loti*, *EXPA1* expression was primarily associated to cells colonized by rhizobia (Figure 1B). Despite the presence of numerous Expansin members in *Lotus*, the available expression data suggest, that *EXPA1* is the isoform mainly recruited both in the intra- and intercellular rhizobial infection. Based on this evidence, we decided to focus on the analysis of *EXPA1* during the two modalities of rhizobial colonization in *Lotus*.

### *EXPA1* promoter activity is linked to rhizobial colonization and nodule organogenesis

To examine in more detail the spatial-temporal expression of *EXPA1* by confocal microscopy, a 2 kb promoter sequence of this gene was fused to a nuclear-localized triple YFP (*pEXPA1::tYFP-nls*). In uninoculated conditions, the *EXPA1* promoter was strongly expressed at the root apex and emerging lateral root primordia, but it was also occasionally detected in the elongation and, differentiation zone as well as in root hairs (Figure 2A-C). In response to *M. loti* R7A inoculation, the *EXPA1* promoter was expressed in deformed root hairs and neighboring epidermal cells, before the initiation of rhizobial infection (Figure 2D). This expression pattern was maintained during IT progression (Figure 2E). Similarly, the promoter activity of *EXPA1* was detected at early steps of the intercellular colonization by IRBG74, such as in twisted root hairs with infection pockets (IP) and in surrounding epidermal cells (Figure 2F). These observations confirm that *EXPA1* is linked to intra- and intercellular colonization events in *Lotus*. However, *EXPA1* also appears to be involved in the nodule organogenesis program, since a prominent promoter expression was found in dividing cortical cells during nodule development (Figure 2G and H).

**Figure 2.**
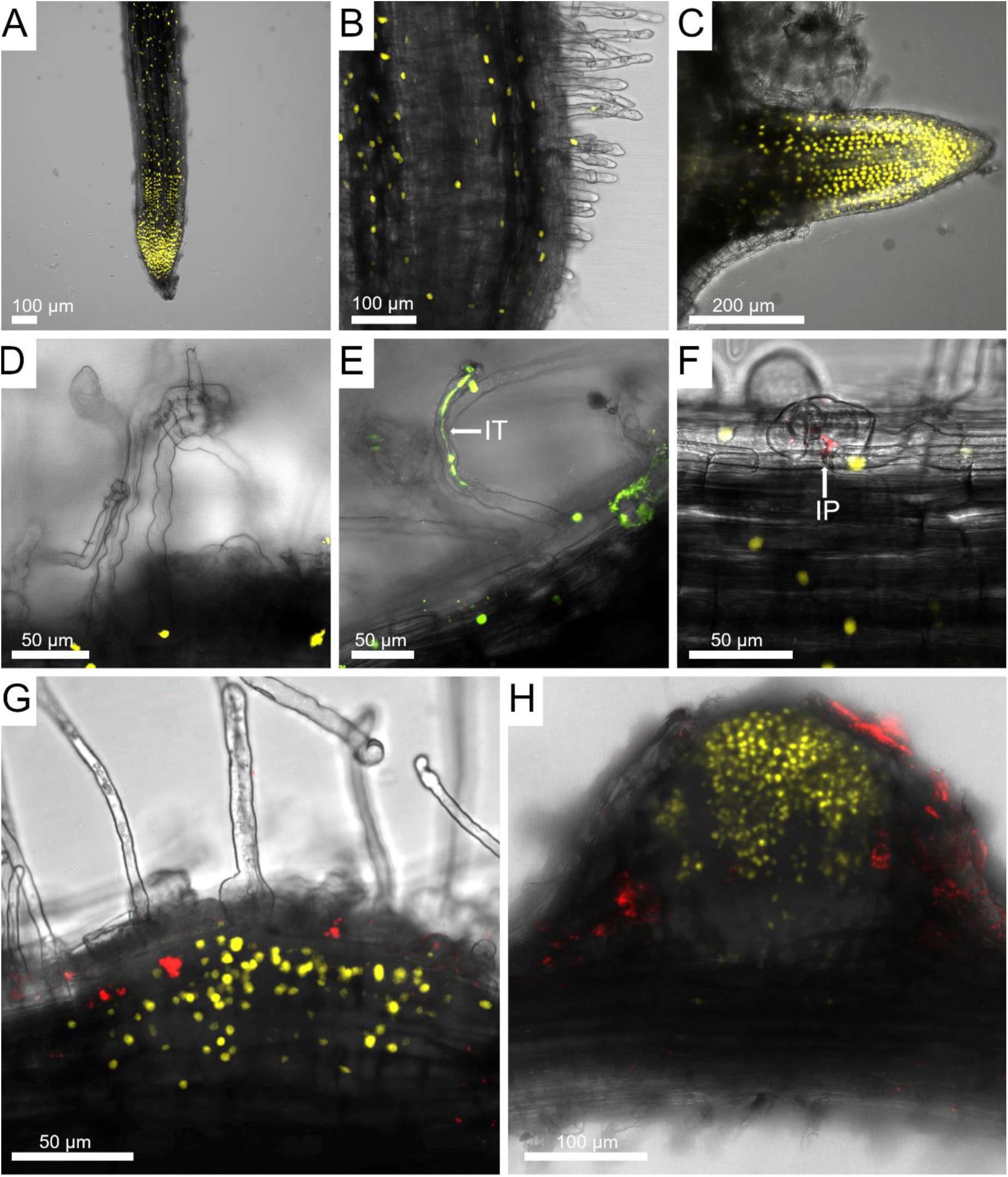
Promoter activity of *LjEXPA1* during intercellular and intracellular rhizobial infection. Confocal microscopy images of *Lotus* transgenic roots expressing the *pLjEXPA1*::*tYFP*-nls construct in uninoculated plants (**A-C**), and after inoculation with *M. loti*-DsRed **(D and E)** and IRBG74-DsRed **(F and H)**. IT, infection thread; IP, infection pocket.

### Subcellular localization of EXPA1 during intra- and intercellular rhizobial colonization

The spatial and temporal expression profile of *EXPA1* place it at critical steps and tissues during the *Lotus*-rhizobia symbiosis. These observations prompted us to explore the localization of this Expansin in *Lotus* roots during rhizobial colonization and nodule development. For this purpose, a construct was generated to express EXPA1 fused to YFP under the control of its native promoter together, with a constitutive expression of a nuclear-localized DsRed as fluorescent transformation marker: *pEXPA1::EXPA1*-*YFP*_*p35S::DsRed-nls*. During infection of transgenic roots by *M. loti*-DsRed, EXPA1-YFP was enriched in the cell wall surrounding the infection chamber (Figure 3A), whereas its presence was barely detectable in the IT cell wall (Figure 3B). Interestingly, a very intense EXPA1-YFP accumulation was also observed at the transcellular passage cleft (TPC), where the IT progresses from cell to cell (Figure 3C).

**Figure 3.**
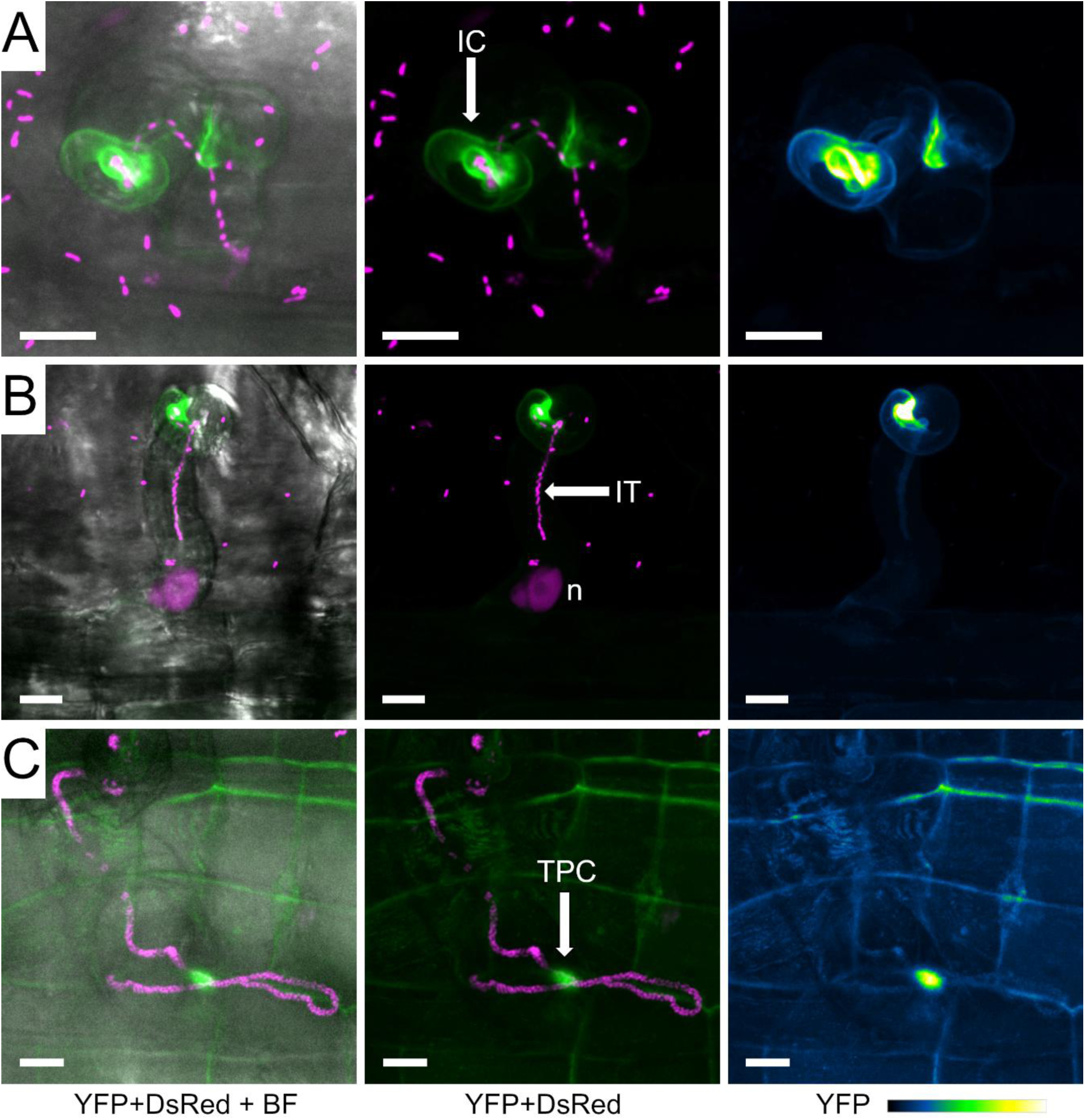
Subcellular localization of LjEXPA1 during intracellular rhizobial infection. Live-cell confocal images of Lotus transgenic roots expressing the *pEXPA1::EXPA1-YFP_p35S::DsRed-nls* construct at different stages of *M. loti* colonization: **A,** Infection chamber (IC) formation; **B,** Infection thread (IT) progression; **C,** Transcellular passage cleft (TPC). Images are maximum intensity projections of z-stacks (step size = 1 μm) showing either the merge of the YFP (green) and DsRed (magenta) channels with (left) or without (middle) the bright field (BF) channel, or the isolated YFP channel (green fire blue, right). At least twelve composite plants from two independent experiments were analyzed. IC = infection chamber; IT = infection thread; TPC = transcellular passage cleft. Scale = 10 μm.

In roots inoculated with IRBG74-DsRed, EXPA1-YFP was detected in specific zones of the cell wall, of twisted root hairs containing infection pockets (IP), a structure that precedes the intercellular colonization of IRBG74 (Figure 4A-C). Additionally, EXPA1-YFP was abundantly recruited at the cell wall of dividing cortical cells during nodule primordia formation after both *M. loti* R7A-DsRed and IRBG74-DsRed inoculation (Figure S3). These results further confirm the relevant presence of EXPA1 at key steps and cellular domains during the intra- and intercellular invasion of *Lotus* roots by *M. loti* and IRBG74, respectively.

**Figure 4.**
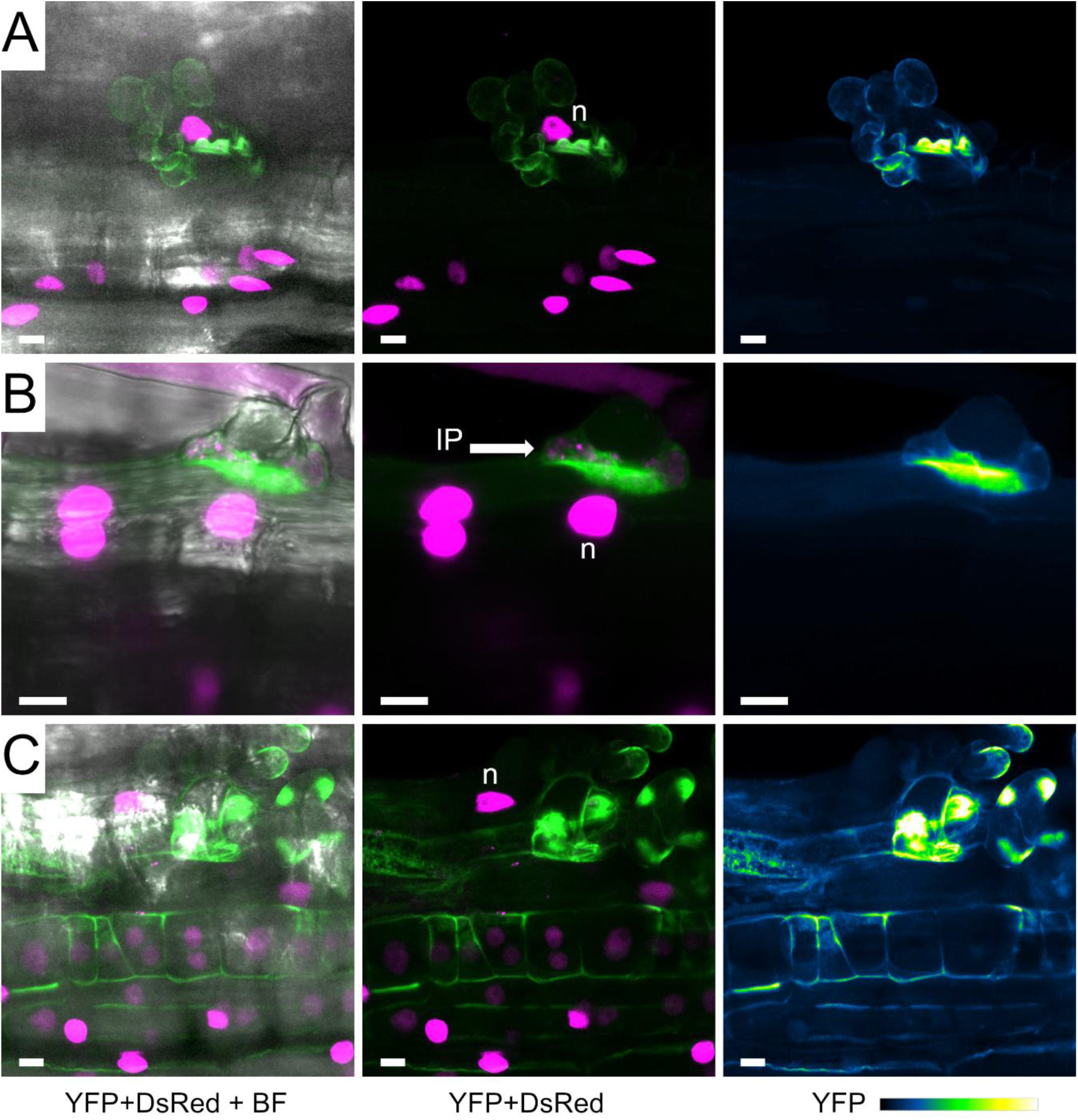
Subcellular localization of LjEXPA1 during intercellular infection. Live-cell confocal images of *Lotus* transgenic roots expressing the *pEXPA1::EXPA1*-*YFP*_*p35S::tDsRed-nls* construct at different steps of intercellular colonization by *A. pusense* IRBG74-DsRed: **A** and **C**, Root hairs swelling and twisting; **B** and **C**, Infection pocket formation (IP). Images are maximum intensity projections of z-stacks (step size = 1 μm) showing the merge of the YFP (green) and DsRed (magenta) channels with (left) or without (right) the bright field (BF) channel. At least twelve composite plants from two independent experiments were analyzed. IC = infection chamber; IT = infection thread; TPC = transcellular passage cleft. Scale = 10 μm.

### Compromised intra- and intercellular symbiotic programs in *expA1* mutants

To determine the relevance of *LjEXPA1* in the *Lotus*-rhizobia symbiosis, two homozygous mutant lines were obtained from the *Lotus* retrotransposon (*LORE1*) mutant collection (Mun *et al*., 2016). *expA1*-1 and *expA1*-2 lines had retrotransposon insertions in the last exon and the 3’ UTR of *EXPA1*, respectively, leading to reduced transcript levels (Figure S4). First, we evaluated the nodulation capacity of the mutants upon inoculation with *M. loti* R7A, by recording their nodulation kinetics. While on wild type Gifu roots (hereafter referred as Gifu), nodule primordia were observed at 1 wpi with *M. loti* R7A, these structures only appeared at 2 wpi in the mutants (Figure 5A). Besides the delay in nodule primordia formation, the nodule numbers were significantly lower at 3-6 wpi in the mutants compared to Gifu wild type plants (Figure 5A). Importantly, the nodules developed by the mutants were smaller and paler, respect to Gifu (Figure 5B-D). Similarly, both size and nodule numbers were affected in *expA1*-1 and *expA1*-2 mutants inoculated with IRBG74 (Figure 5E-H).

**Figure 5.**
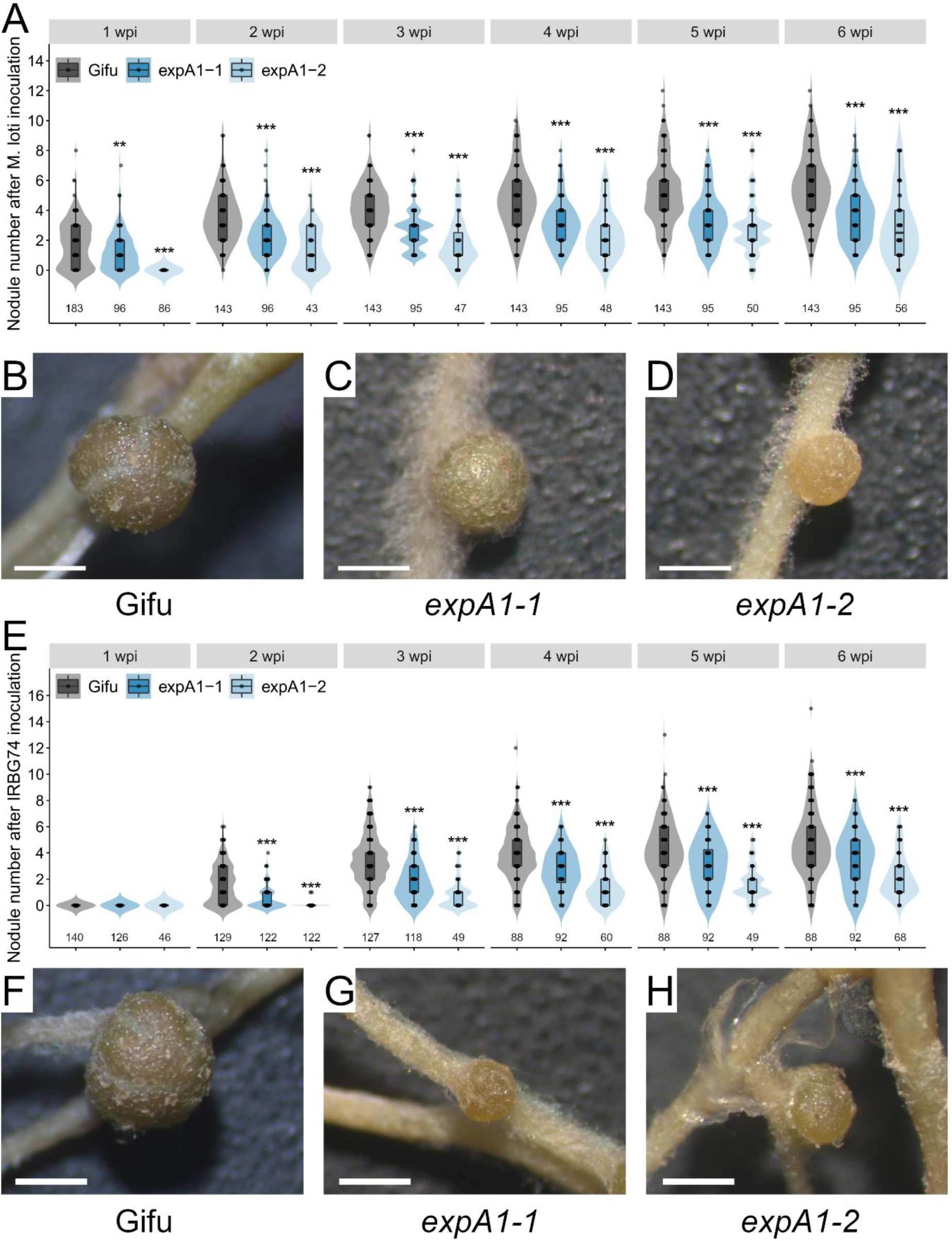
Compromised nodulation of *expA1* and *expA2* mutants after inoculation with *M. loti* R7A and IRBG74. Total number of nodules in Gifu (n=20), e*xpA1-1* (n=20) and e*xpA1-2* (n=20) at 1-6 wpi with *M. loti* **(A)** and IRBG74 **(E)**. Representative images of 3 weeks old nodules formed in Gifu **(B and F)**, *expA1-1* **(C and G)** and *expA1-2* **(D and H)** at 6 wpi with *M. loti* **(B-D)** and IRBG74 **(F-H)**. Scale 1 cm. In boxplots, the center line represents means values of 3 independent experiments; box limits, upper and lower quartiles; whiskers, 1.5× interquartile range; points represent individual data points. The asterisk indicates statistical significance between the *LORE1* mutant and Gifu according to Mann–Whitney U-test (\**P* < 0.05; \*\**P* < 0.01; \*\*\**P* < 0.001).

The collected evidence prompted us to monitor the two modalities of rhizobial colonization in the two *expansin* mutants. In Gifu plants, the typical intracellular infection through root hair ITs was observed at 7 dpi with *M. loti* R7A (Figure 6A). By contrast, at this timepoint ITs were not observed in the *expA1*-1 and *expA1*-2 mutants, and rhizobia were rather found attached to the root hair tip or within infection chambers (Figure 6B and C). Quantitative analysis of these observations indicated that epidermal and cortical ITs *per* root were reduced by over 90 % in *expA1*-1 and *expA1*-2 mutants compared to Gifu (Figure 6G). Subsequently, we evaluated the intercellular colonization of *A. pusense* IRBG74 in the different genotypes, following the approaches that we have established to assess the *Lotus*-IRBG74 symbiosis (Montiel et al. 2021). In accordance with previous reports, IRBG74 promotes the formation of infection pockets in swollen and twisted root hairs at 7 dpi (Figure 6D). These responses also occurred in the *expA1*-1 and *expA1*-2 mutants inoculated with *A. pusense* IRBG74, but apparently with fewer and less colonized infection pockets (Figure 6E and F). Based on these findings, we followed a root endosphere protocol to evaluate by qPCR, the rhizobial *nodA* abundance in genomic DNA samples isolated from roots of Gifu and the *Expansin* mutants at 3 wpi with IRBG74. The presence of the IRBG74-*nodA* gene was 64-87% lower in the *expA1*-1, and *expA1-*2 mutants compared to Gifu, reflecting a compromised intercellular colonization of the roots by IRBG74 (Figure 6H).

**Figure 6.**
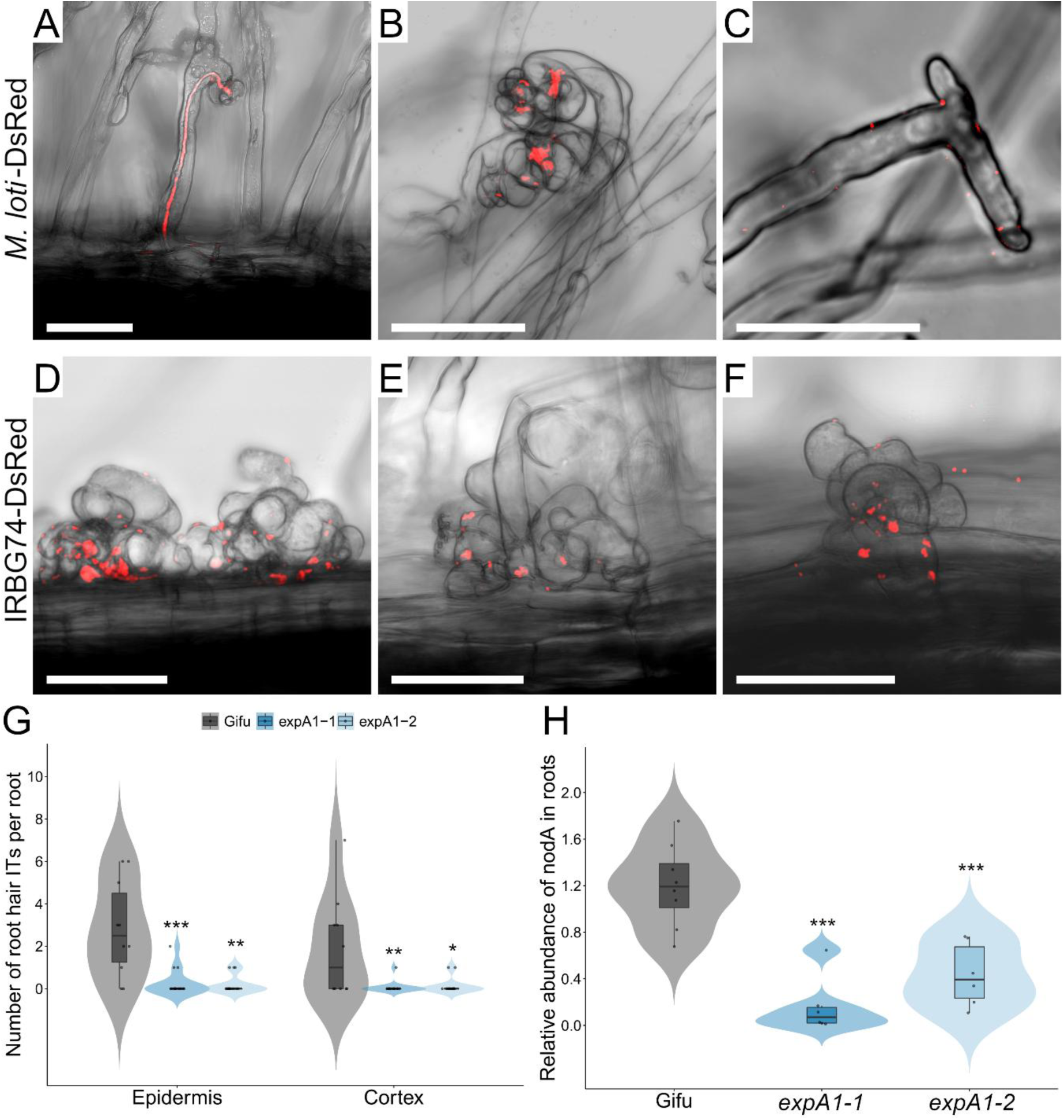
Altered rhizobial infection in *expA1* mutants. Confocal microscopy images of rhizobial colonization in Gifu (**A** and **D**), *expA1*-1 (**B** and **E**) and *expA1*-2 (**C** and **F**) roots at 7 dpi with *M. loti*-DsRed (**A**-**C**) and IRBG74-DsRed (**D**-**F**). **G**, Number of root hair ITs found in the epidermis and cortex at 10 dpi with *M. loti*–DsRed on Gifu (n = 10), *expA1*-1 (n = 15) and *expA1*-2 (n = 15). **H**, Abundance of IRBG74-*nodA* by qPCR in genomic DNA isolated from Gifu (n = 8), *expA1*-1 (n = 6) and *expA1*-2 (n = 6) at 3 wpi with IRBG74 and normalized to the *LjNFR5* gene accumulation.

## Discussion

### Differential expression of *Lotus* Expansins in biotic interactions

Over recent decades, a range of studies have documented that Expansins form a multigene family in plants, mostly composed by four subfamilies (Cosgrove, 2024, Wang *et al*., 2024). The selected species in our phylogenetic analysis show a higher abundance of the alpha subfamily with an increased representation of the EXLB group in legumes relative to non-legumes, an observation that is consistent with other studies (Guimaraes *et al*., 2017, Wang *et al*., 2024). While the alpha and beta subfamilies are extensively documented in the literature for loosening the plant cell wall during developmental programs and in response to (a)biotic stimuli, reports on the roles of EXLA and EXLB members remain limited (Cosgrove, 2015). When analyzing the transcriptomic profile of *Lotus* Expansins in response to biotic interactions, we found that most of the *EXLBs* increase their expression after inoculation with the pathogens *Pseudomonas syringae* and *Ralstonia solanacearum* (Kelly *et al*., 2018). Different studies conducted in *Arachis* spp., confirm the relevant role of *EXLB* in response to pathogen perception, since the heterologous ectopic expression of *Arachis duranensis EXLB8* confers resistance towards *Sclerotinia sclerotiorum* and *Meloidogyne incognita* attack (Guimaraes *et al*., 2017, Brasileiro *et al*., 2021). Herein, we observed that expression of one *EXLB* gene (LotjaGi4g1v0195900) was significantly induced at 1 dpi with *M. loti*. The analysis of the expression data in *Lotus* also revealed that during arbuscular mycorrhiza symbiosis (AMS), Expansins of the four subfamilies are notably upregulated. In contrast, only a small subset of Expansins exhibit high expression levels in nodules or at initial stages of intra/intercellular rhizobial colonization. Among these, LotjaGi1g1v606600, which encodes an EXPB isoform, is the potential ortholog of soybean EXPB1, based on its phylogenetic distribution. *GmEXPB1* promoter is strongly expressed in different tissues of developing soybean nodules, and transgenic plants with altered transcript levels of *GmEXPB1* by RNAi or overexpression, are affected in nodule size and number (Li *et al*., 2015).

Likewise, the involvement of alpha Expansins in various biotic interactions has been documented. In *Nicotiana benthamiana* leaves, Turnip mosaic virus (TuMV) leads to a downregulation of both transcript and protein levels of the plasmodesmata-specific EXPA1. Furthermore, overexpression of *NbEXPA1* increases resistance to TuMV infection, while its silencing decreases it (Park *et al*., 2017). Conversely, in *Nicotiana tabacum*, overexpression of the alpha Expansin *EXPA4* promotes disease progression caused by *Pseudomonas syringae* DC3000 (Chen *et al*., 2018). While several studies show that at different stages of the legume-rhizobia symbiosis different *Expansins* are transcriptionally induced (Gyorgyey *et al*., 2000, Giordano and Hirsch, 2004, Libourel *et al*., 2023), their specific functional roles remain scarcely characterized. In *Lotus*, six alpha Expansins were significantly upregulated at 1-3 dpi with *M. loti*, whereas only four members of this subfamily were induced by IRBG74 at 3-10 dpi. Notably, the expression of 2 alpha Expansins was repressed in response to *M. loti*, and one after IRBG74 inoculation, suggesting that certain Expansins might play a negative role during rhizobial infection.

### Cell wall rearrangement during rhizobial colonization and nodule development

Prior to the formation of the IT, the microsymbiont attaches to the apical region of the root hair, leading to swelling and curling of the root hair, that traps the rhizobia within an infection chamber. This process requires significant cellular changes, as the apical growth of the root hair cell wall is reversed (Fournier *et al*., 2008, Gao *et al*., 2024). Recent evidence revealed that Glycoside Hydrolase 9C2, a cell wall degrading enzyme, is abundantly accumulated in the infection chamber (IC) of *M. truncatula* root hairs. In addition, mutant lines disrupted in *GH9C2* display aberrant ITs (Zhao *et al*., 2025). Similarly, abnormal IC are formed in the *nodulation pectate lyase* (*npl*) mutant in *L. japonicus* (Xie *et al*., 2012). These studies reveal that enzymatic degradation of cell wall components in root hairs, mediated by GH9C and NPL, is essential for intracellular rhizobial infection. Our study presents evidence that also EXPA1 contributes significantly to intracellular symbiotic colonization in *L. japonicus*. EXPA1 was prominently localized within the IC, and mutants deficient in *EXPA1* showed marked defects in IT development. Importantly, EXPA1 strongly accumulated at the so-called transcellular passage cleft (Zhang and Ott, 2024). This localization pattern correlates with the cytolocalization of NPL and symbiosis-specific pectin methyl esterase (SyPME1) in *M. truncatula*, where the coordinated action of both proteins is essential to alter the IT cell wall biomechanics during the transcellular invasion of rhizobia (Su *et al*., 2023). Our results show that, in addition to the enzymatic activity of cell wall remodeling proteins, the non-catalytic cell wall relaxation mediated by EXPA1, occurs at critical stages of intracellular rhizobial infection.

The molecular mechanisms underlying intercellular colonization in legume-rhizobia symbiosis remain largely unclear, even though this process occurs in roughly 25% of nodulating legume genera (Sprent 2007). In this regard, we recently reported that *A. pusense* IRBG74 infects *Lotus* roots intercellularly, establishing it as a novel working model to study this process (Montiel *et al*., 2021). Notably, both the total number of nodules and the number of pink nodules is comparable in plants inoculated with either IRBG74 or the cognate rhizobial strain *M. loti*. In contrast, the *Ljnlp* mutant exhibits a more pronounced reduction in nodule formation within the *Lotus*-IRBG74 symbiosis than in the *Lotus*-*M. loti* association (Montiel et al. 2021), indicating a greater impact on the intercellular infection mechanism by the loss of function in the symbiotic pectate lyase. An initial response observed during the intercellular colonization of IRBG74 in *Lotus* roots is extensive curling and twisting of root hairs. This event is subsequently followed by the formation of infection pockets (IP) within the root hairs; from these IP, the bacteria migrate intercellularly toward a deeper layer of the root tissue (Montiel *et al*., 2021, Quilbe *et al*., 2022). At these steps of intercellular infection, EXPA1 was particularly abundant at the periphery of swollen root hairs and IP, suggesting that cell wall remodeling by EXPA1 takes place in these processes.

EXPA1 was prominently expressed in dividing cortical cells during nodule primordia development in plants with either intercellular or intracellular colonization. Furthermore, the reduction in nodule formation observed in *expA1* mutants was comparable when inoculated with either *M. loti* or IRBG74. While intracellular and intercellular infections involve distinct transcriptomic and cellular pathways, a set of genes is fundamental for both processes (Garcia-Soto *et al*., 2023, Montiel *et al*., 2023). Our findings indicate that *EXPA1* represents one such gene.

In summary, structural remodeling of the cell wall matrix at confined cellular domains in legume roots is crucial for rhizobial colonization, with different cell wall remodeling proteins being synergistically involved at multiple steps. Recent research has provided substantial insight into the intricate interplay among these components, but the picture is far from being complete. In this frame, the additional identification of members belonging to other well-known cell wall remodeling protein families such as the Expansins and the characterization of their role during symbiotic infection constitute a relevant contribution to the field.

## Supplementary information

**Figure S1. Variable expression pattern of *Lotus Expansins* in different organs and tissues.** Heat map gene expression of *Lotus Expansins* in aerial and ground tissues/organs, under non-symbiotic, symbiotic and pathogenic conditions. Data collected from *Lotus* base (https://lotus.au.dk/). AMS, arbuscular mycorrhiza symbiosis; RNS, root nodule symbiosis.

**Figure S2. Phylogeny and composition of the Expansin family in different plant species**. Phylogenetic distribution **(A)** and number **(B)** of alpha, beta, alpha-like and beta-like Expansins in *Physcomitrella patens, A. thaliana, Oryza sativa, Glycine soja, Phaseolus vulgaris, Medicago truncatula* and *L. japonicus*. EXPA1 is highlighted with an arrow. LotjaGi1g1v060600, marked with an asterisk, is the putative orthologue of GmEXPB2 (Li et al. 2015). **C)** Identity matrix of *Lotus* Expansins represented by heatmap using the predicted amino acid sequences. Made on Uniprot (https://www.uniprot.org/align/).

**Figure S3. Subcellular localization of LjEXPA1 during nodule primordia formation.** Live-cell confocal images of *Lotus* transgenic roots expressing the *pEXPA1::EXPA1*-*YFP*_*p35S::tDsRed-nls* construct during nodule primordia formation after *M. loti* R7A-DsRed **(A)** and IRBG74-DsRed colonization **(B)**. Images are single focal planes **(A)** or maximum intensity projections of z-stacks (**B**, step size = 1 μm) showing either the merge of the YFP (green) and DsRed (magenta) channels with (left) or without (middle) the bright field (BF) channel, or the isolated YFP channel (green fire blue, right). At least twelve composite plants from two independent experiments were analysed. Scale = 50 μm.

**Figure S4. Impact on transcript levels of *LjEXPA1* by *LORE1* insertions. A,** Schematic representation of *LjEXPA1* gene disrupted by LORE1 insertions. Black boxes: exons; Dashed lines, UTRs; Solid lines, introns. Retrotransposon insertions are indicated with red triangles. **B,** Boxplots represent the normalized expression levels of *EXPA1,* calculated by RT-qPCR from 4 independent biological replicates (n = 5 per biological replicate) in Gifu, *expa1*-1 and *expA1*-2 roots.

## Author contributions

J.M. and J.S. planned and designed the research. J.M., I.G.S, B.L., E.M.M., N.S. and M.V., performed experiments. J.M., B.L., T.O. and J.S. revised the manuscript and provided resources and infrastructure. J.M. and E.M.M. wrote the manuscript.

## Acknowledgements

We thank Mariana Robledo Gamboa for identifying the Expansin sequences in different plant species.

