## Supplementary figures and images for "The *Lotus japonicus* alpha Expansin *EXPA1* is recruited during intracellular and intercellular rhizobial colonization"

### Figure S1

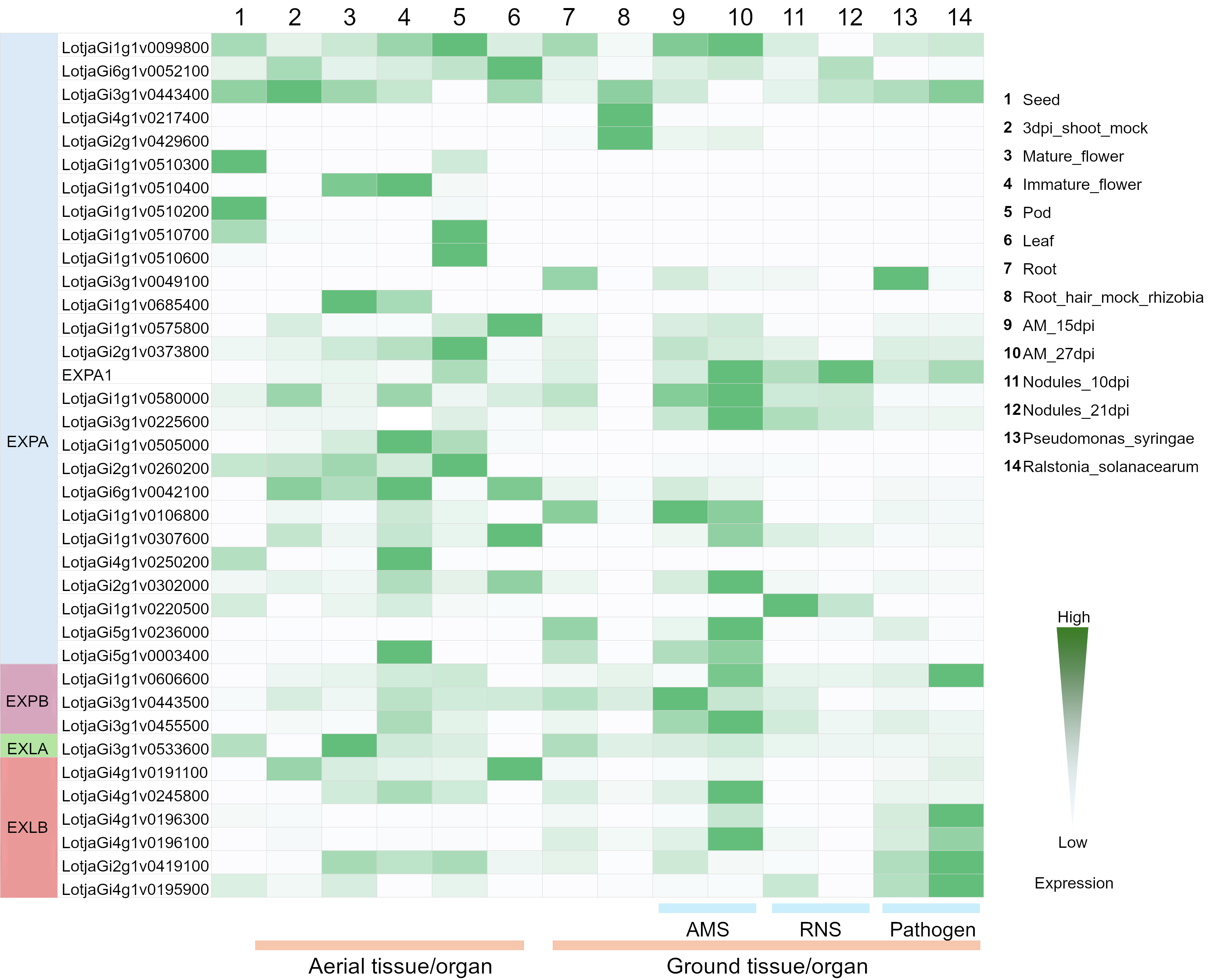

### Figure S2

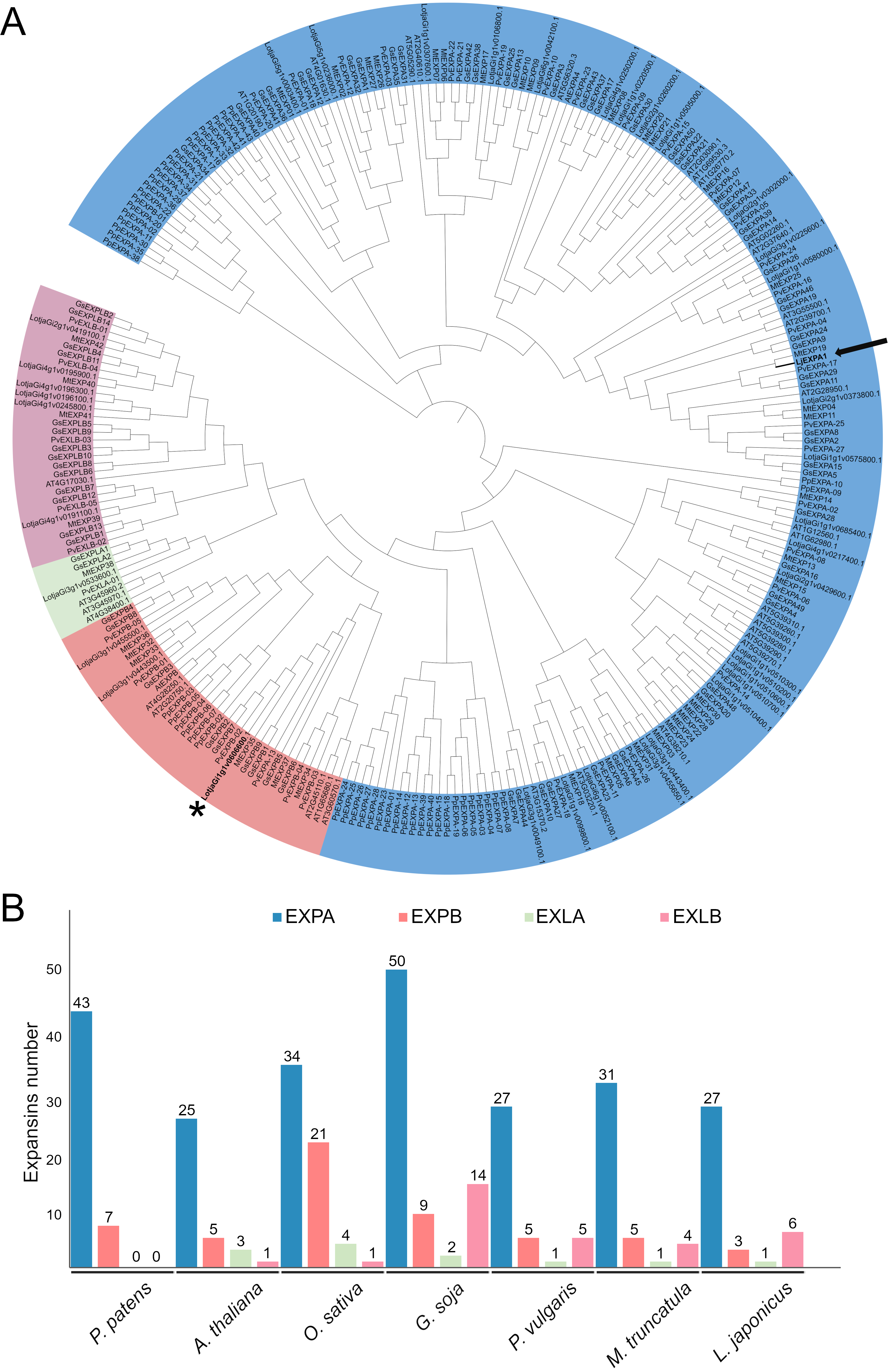

### Figure S3

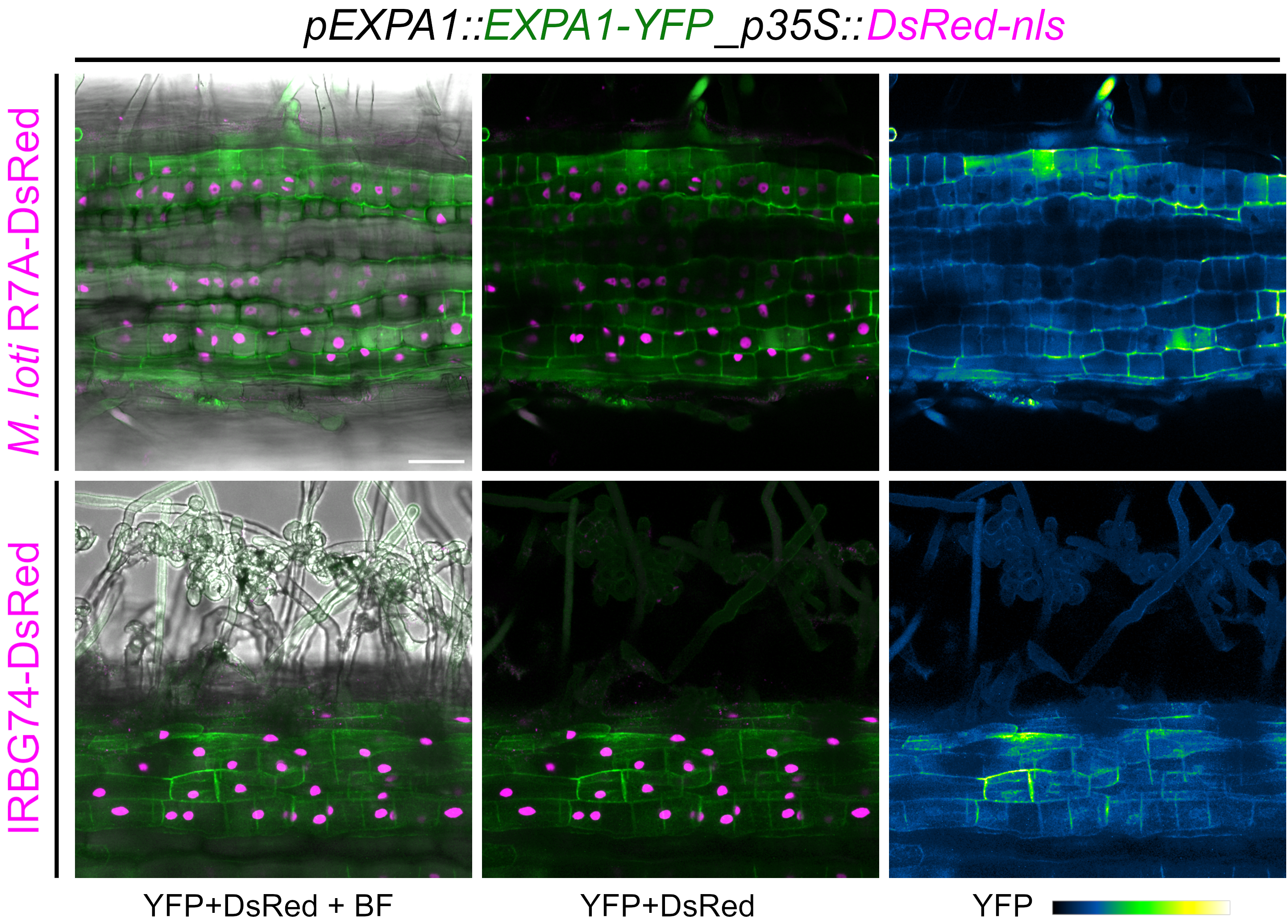

### Figure S4

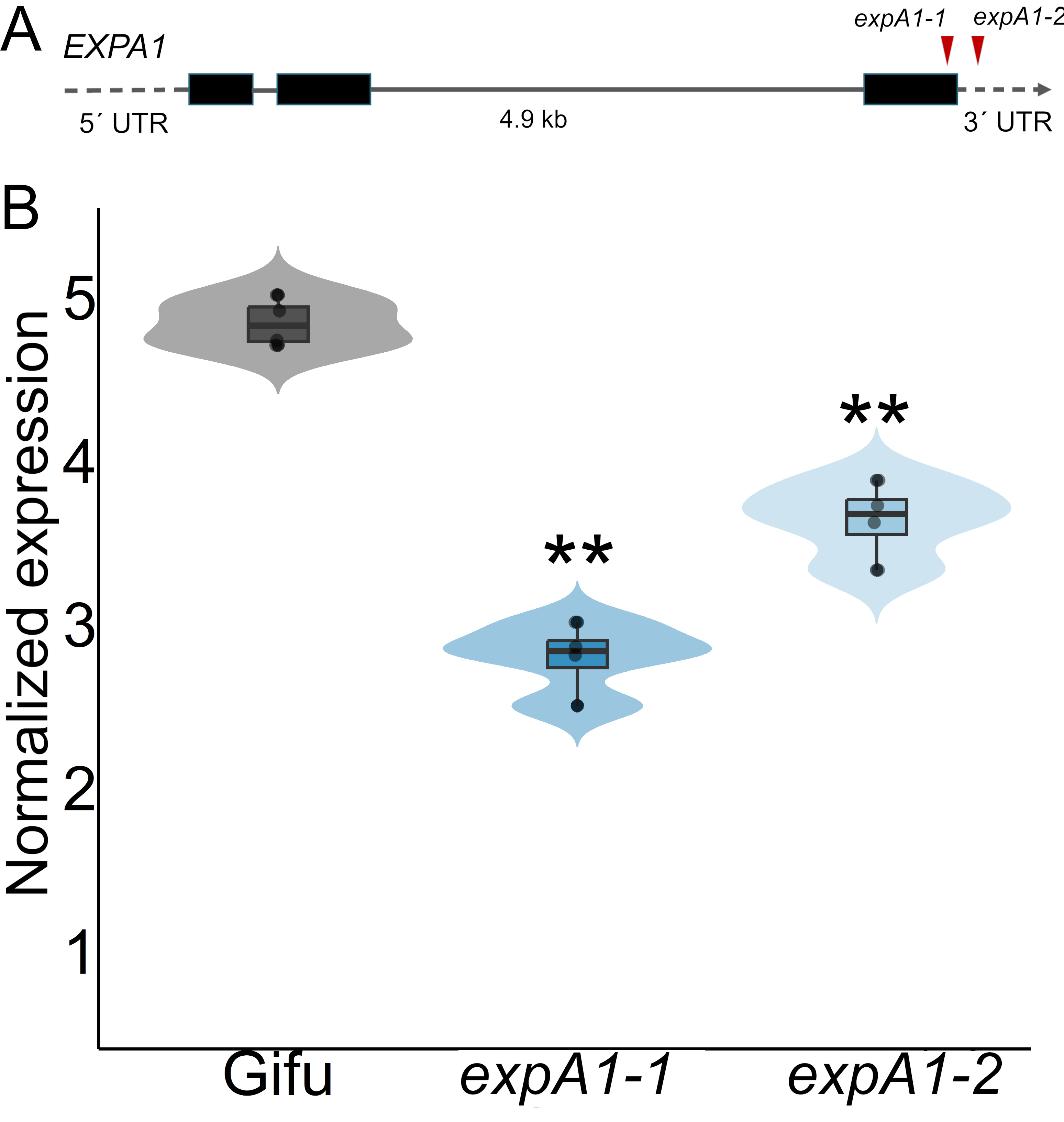
